# Oligomeric α-Synuclein induces skin degeneration in a reconstructed human epidermis model

**DOI:** 10.1101/2021.06.17.448863

**Authors:** Júlia T. Oliveira, Rodrigo De Vecchi, Vanja Dakic, Gabriela Vitória, Carolina Pedrosa, Mayara Mendes, Luiz Guilherme H.S. Aragão, Thyago R. Cardim-Pires, Daniel Rodrigues Furtado, Roberta O. Pinheiro, Débora Foguel, Lionel Breton, Charbel Bouez, Marilia Zaluar P. Guimarães, Stevens Rehen

## Abstract

Cell senescence may promote epidermal inflammation and degeneration, termed as inflammaging, which is accompanied by keratinocyte loss, resulting in fine lines of wrinkles. Recent findings showed that healthy elderly skin expresses age- and neuron-related amyloidogenic proteins, such as tau, β-Amyloid34, and α-synuclein (α-Syn), typically found in patients with neurodegenerative diseases. These proteins form toxic aggregates that trigger inflammatory signals. Herein, we investigated the impact of oligomeric α-Syn (Oα-Syn) on the neurosphere (NP) and the reconstructed human epidermis (RHE) 3D models. First, we found the expression of α-Syn, β-Amyloid, and amyloid precursor protein (APP) in the RHE. Second, we challenged the RHE and NP with Oα-Syn, which decreased RHE regeneration, measured by the percentage of cell proliferation and thickness of the stratum basale, but did not affect NP neurite outgrowth. Oα-Syn did not decrease the number of human neonatal epidermal keratinocytes (HEKn) but, as seen for the RHE, it also decreased the proliferation of HEKn. We confirmed that the oligomeric, and not the monomeric α-Syn species, accounted for the proliferation-decreasing effect. Oα-Syn also increased the NF-kB nuclear translocation in HEKn analyzed by nucleus/cytoplasm NF-κB fluorescence intensity. In addition, Oα-Syn triggered inflammation in the RHE, by increasing the mRNA levels of IL-1β and tumor necrosis factor-alpha (TNF-α), and the release of TNF-α in a time-dependent manner. These findings show that Oα-Syn does not affect neurite outgrowth but induces a decrease in keratinocyte proliferation along with epidermal inflammation. With our tridimensional models, we demonstrated that the neurodegenerative protein Oα-Syn also degenerates the epidermis, drawing attention to the need of target-based screening to prevent and treat the effects of skin aging.

## Introduction

The epidermis is the outermost layer of the skin and is mainly composed of keratinocytes. The proliferation of keratinocytes is restricted to the basal layer, which is crucial to skin homeostasis and regeneration. Chronological skin aging, which is accompanied by epidermal degeneration, is a natural process throughout life. However, it can be accelerated by some factors, such as smoking (Martires et al., 2009) and ultraviolet (UV) exposure (Baumann, 2007). Aged skin displays high levels of inflammatory mediators including the proinflammatory cytokine tumor necrosis factor-alpha (TNF-α). This cytokine triggers inhibition of DNA repair during G2/M checkpoint and apoptosis in keratinocytes (Faurschou, 2010; Borg et al., 2013). High levels of TNF-α are also found in aged skin as a compensatory mechanism in response to permeability barrier dysfunction (Hu et al., 2017). TNF-α prompts the translocation of the nuclear factor kappa-light-chain-enhancer of activated B cells (NF-κB) into the nucleus, leading to the transcription of additional inflammatory genes (Pupe et al., 2003; Lan et al., 2005). NF-κB plays a pivotal role in skin homeostasis and aging (Grinberg-Bleyer et al., 2015). Aggregation of amyloidogenic proteins in the central nervous system, such as α-Synuclein (α-Syn) and β-Amyloid (Aβ), are the pathological hallmarks of neurodegenerative processes underlying Parkinson’s and Alzheimer’s diseases, respectively. The leading molecular signature in Parkinson’s disease is the accumulation of misfolded α-Syn, including oligomeric α-Syn (Oα-Syn), in dopaminergic neurons in the substantia nigra. However, a growing body of evidence demonstrates abnormal aggregates in other tissues with consequences to physiology and pathology (Wakabayashi et al., 1990; Beach et al., 2010; Gelpi et al., 2014; Braak and Del Tredici, 2017). For instance, neurodegenerative disease-related proteins, such as tau, Aβ34, and α-Syn, were found in the epidermal layer of human skin samples (Rodriguez-Leyva et al., 2017; Akerman et al., 2019b). In the case of Parkinson’s disease patients, α-Syn substantially accumulates in the epidermis — also occurring in healthy individuals to a lesser extent (Rodriguez-Leyva et al., 2016). Although studies have detected α-Syn and other amyloidogenic proteins in human skin, the knowledge on how they affect skin homeostasis is not clear yet. Evidence from studies in the nervous system suggest inflammation among the consequences of protein misfolding (Alvarez-Erviti et al., 2011). As the epidermis is composed of peripheral nerve endings, in this work, we investigated the impact of Oα-Syn on neurons and epidermis 3D models, termed neurospheres (NP), and reconstructed human epidermis (RHE), respectively. We found that Oα-Syn did not affect neurite outgrowth but it decreased keratinocyte proliferation, consequently decreasing epidermal thickness. Additionally, Oα-Syn elicited keratinocyte inflammation by increasing NF-κB nuclear translocation and cytokine production. Taken together, our results highlight a novel toxic role of Oα-Syn in the human epidermis that may contribute to skin degeneration underlying aging.

## Materials and Methods

### Cell culture

#### Human Epidermal Keratinocytes, neonatal (HEKn)

Human neonatal epidermal keratinocytesl (HEKn) (Life Technologies - C0015C) were cultivated in EpiLife™ Medium (Thermo Fisher Scientific - MEPI500CA) with 1x HKGS (Thermo Fisher Scientific - S-001-5) on a T75 culture flask. When HEKn achieved 80% of confluence, they were split with TrypLE™ Express Enzyme (1X) (Thermo Fisher Scientific - 12605028) for 10 minutes, counted, and centrifuged at 200 x *g* for 10 minutes. Then, the cells were replated at 3 x 10^5^ cells per T75 culture flask or at 1.2 x 10^4^ cells per well in 96-well plates and maintained in an incubator at 37 °C with 5% CO2. The medium was changed every other day.

#### Reconstructed Human Epidermis (RHE)

SkinEthic™ RHE was reconstructed *in vitro* from healthy human keratinocytes, which were donated by volunteers with signed consent terms, and expanded before use. RHE was grown on an inert polycarbonate filter (0.5 cm^2^) at the air-liquid interface, in a chemically defined medium (Rosdy and Clauss, 1990). RHE is a model of a multilayered human epidermis, with clearly visible basal, spinous, granular, and corneal layers. This model features a functional permeability barrier and is reliable to test skin permeability and toxicity (Tornier et al., 2010). Each RHE batch was checked according to standard quality control criteria, and the tests included viability, barrier function, and morphology (De Vecchi et al., 2018). Seventeen-day-old RHE was used in the experiments because it is the time point when RHE achieves maturity, meaning it has all layers completely formed. All substances were diluted in the medium, so administered from the stratum basale side.

#### Neurosphere (NP)

The method of neural stem cells generation is based on a protocol developed by Life Technologies (# MAN0008031) (Yan et al., 2013). We cultivated and split NSC on neural expansion medium (NEM) (Advanced DMEM/F12, Thermo Fisher Scientific - 12634-010, and Neurobasal medium, Thermo Fisher Scientific-12348017, 1:1, plus 1% neural induction supplement, Thermo Fisher Scientific - A1647701) on a surface previously coated with Geltrex LDEV-Free (Thermo Fisher Scientific, A1413302). When neural stem cells culture achieved 90% of confluence, they were split with Accutase (Merck Millipore, SCR005), counted, and 9 x 10^3^ cells in 150 μL NEM were plated per well of a round bottom Ultra-low attachment 96-well plate (Corning-Costar, ref: 7007). Then, we centrifuged the plate at 300 x *g* for 5 minutes and maintained it in an incubator at 37 °C with 5% CO2. Forty-eight hours later, we removed 100 μL of medium and added 150 μL of differentiation medium: DMEM-F12, Life Technologies - 11330-032, Neurobasal medium, 1X N2, Invitrogen −17502001, 1X B27, Thermo Fisher Scientific- 17504001. The medium was replaced every 2-3 days until day 10 when each NP was individually plated for allowing neurite outgrowth on surfaces previously coated with 100 μg/mL poly-L-ornithine (Sigma Aldrich, P3655) and 20 μg/mL laminin (Invitrogen, 23017-015) with conditioned medium and fresh differentiation medium (1:1). Twenty-four hours after plating, NPs were challenged with 30 μM Oα-syn and, 24 hours later, fixed for immunostaining procedures.

### Oligomers preparation

#### α-Synuclein (α-Syn)

The wild-type α-Syn monomers were expressed and purified according to Kruger et al., 1998 (Krüger et al., 1998). We added an additional step to the purification procedure to remove LPS, a bacterial endotoxin, that could interfere with the results in cell culture assays. To obtain a LPS-free protein, purified α-Syn monomers were passed through a polymyxin B-conjugated resin (Detoxi-gel endotoxin removing column, Thermo Scientific, Waltham, MA, USA). α-Syn oligomers (Oα-Syn) were produced by incubating 120 μM monomers in PBS for 18 h at pH 7.5, 37 °C under stirring (800 RPM) in a thermomixer. The presence of oligomers was confirmed by TEM imaging (not shown).

#### Aβ peptide

We added 1 mg Aβ peptide (1-42) (Anaspec, CA, USA) into a microtube containing 270 μL of 1,1,1,3,3,3-Hexafluoro-2-propanol (HFIP) and left it diluting for 1 hour in the hood without homogenization. We transferred the microtube to the ice for 5-10 minutes, aliquoted (60 μL/ 600 mL microtube) it, and left the microtubes open inside the hood overnight to allow the HFIP to evaporate and form a film. The films were resuspended with 10 μL DMSO scraping the lateral of the microtubes using the tip of the pipette. We added 490 μL cold PBS per microtube and homogenized it using a vortex mixer for 10 seconds. We also prepared a vehicle sample (10 μL DMSO plus 490 μL cold PBS). We kept the microtubes at 4 °C for 20-24 hours. Then, we centrifuged them at 14,000 x g for 10 minutes at 4 °C and transferred the supernatant to a low binding microtube. We used 30 μL of the preparation for HPLC characterization and another 30 μL for protein dosage by the BCA method (Pierce BCA protein assay – Thermo Fisher Scientific). Finally, we kept the microtubes at −80 °C until we thawed and used them for the experiments.

### Sample preparation and immunostaining

RHE, HEKn, and NPs were fixed in 4% paraformaldehyde for 20 minutes and washed 3 times in phosphate-buffered saline (PBS). RHE samples were embedded in a Tissue-Tek OCT compound and frozen in liquid nitrogen. Longitudinal cryosections were cut at 10 μm on a cryostat (Leica CM 1850) and mounted on gelatin precoated slides. The immunostained procedure occurred as follows. Samples were permeabilized with 0.3% triton x-100 for 15 minutes, blocked in 3% bovine serum albumin (BSA) for 1 hour at RT, and incubated overnight at 4°C in a solution containing primary antibodies diluted in BSA 2%. Then, we washed samples 3 times in PBS, reblocked in 2% BSA for 20 minutes, and incubated in a solution containing corresponding secondary antibodies diluted in 2% BSA for 40 minutes at RT. Finally, samples were rinsed 3 times in PBS, stained with the nuclear marker DAPI (0.5 μg/mL) for 5 minutes, rinsed again 3 times in PBS, and slides were mounted with Aqua-Poly-mount (Polysciences) or cell plate wells were covered with glycerol (Sigma). We used the following primary antibodies: mouse anti-Ki67 (1:50, 550609, BD Pharmigen); mouse anti-α-synuclein (1:200, AHB0261, Thermo Fisher Scientific); mouse anti-NF-κB p65 (1:200, SC-8008, Santa Cruz Biotechnology); mouse anti-Alzheimer Precursor Protein (APP) 22C11 (1:100, MAB348, Millipore), and mouse anti-β-Tub III/Tuj 1 (1:4000, MO15013, Neuromics). The secondary antibodies used were: Goat anti-Mouse IgG (H+L) Secondary Antibody, Alexa Fluor® 488 conjugate (A-11001, Invitrogen); and Goat anti-Mouse IgG (H+L) Secondary Antibody, Alexa Fluor® 594 conjugate (A-11032, Invitrogen).

The hematoxylin-eosin-safran staining was performed on paraffin sections using the Sakura Tissue-Tek® Prisma® automated slide stainer, according to the manufacturer’s protocol.

### Image acquisition

Images of immunostained RHE slides were acquired with a confocal microscope Leica TCS SP8 using a 63x objective lens, while images of immunostained HEKn and NPs were acquired with the high content screening microscope Operetta (PerkinElmer, Waltham, MA, USA) using a 2x or 40x objective lens, respectively. For capturing HEKn images, 9 fields per well were systematically selected.

### Proliferation and nuclei counting assay

HEKn and RHE were immunostained for the proliferation marker Ki67 as described above. For RHE analysis, we quantified the ratio of total Ki67+ nuclei/ total DAPI+ nuclei and converted it into percentages. All images were processed and quantified using ImageJ software. HEKn analysis was performed by the software Columbus Image Data Storage and Analysis System (PerkinElmer). We selected the total amount of nuclei using the DAPI channel. Then, we found the Ki67+ nuclei using the Alexa 488 channel, which was the emission spectrum of the secondary antibody used, inside the pre-selected DAPI+ regions. Results were defined by the ratio of Ki67+ nuclei/ DAPI+ nuclei and plotted as the percentage of positive cells. Quantification of DAPI-stained HEKn nuclei was used as an indicator of cell survival, and the results were presented as the percentage of DAPI+ nuclei relative to control.

### Neurite outgrowth assay

Plated NPs were immunostained for β-tubulin III, a neuronal cytoskeleton marker. Images were acquired with the high content screening system Operetta, and analyzed using Neuromath software. The analysis was defined by the total number of neurites crossing a circular mask created at 4x the diameter of NP’s core (Supplementary figure 1). Results were obtained from 1 experiment comprising 20 technical replicates for each experimental group.

### NF-κB nuclear translocation assay

NF-kB nuclear translocation assay is based on the amount of nuclear NF-kB relative to its cytoplasmic counterpart, therefore being a predictor of inflammation. We treated HEKn and RHE with 2, 10, or 20 ng/mL TNF-α (TNA-H4211, ACROBIosystems, DE, USA) for 45 minutes, fixed cells with PFA 4% for 30 minutes, and performed immunostaining for NF-κB subunit, p65/RelA. For RHE analysis, we created a mask to identify the cytoplasm immunostained for NF-κB p65 antibody and the nuclei stained with DAPI using the same grayscale and threshold for all images. Immunostaining intensity from the two masks, which was represented as arbitrary units (A.U), was quantified and plotted as the ratio: nucleus/cytoplasm NF-κB fluorescence intensity. All images were processed and quantified using ImageJ software. For HEKn analysis, we systematically selected 9 fields per image and used the high-content image analysis software Harmony 5.1 (PerkinElmer, Waltham, MA, USA) to generate the ratio of nucleus/cytoplasm intensity.

### Quantitative RT-PCR (RT-qPCR)

Reconstructed Human Epidermis was sliced into thick sections, transferred to 2.0 mL pre-filled tubes with 1.5 mm Triple-Pure™ Zirconium homogenizer beads (Benchmark Scientific, USA) containing a fresh amount of Lysis Buffer (ThermoFisher Scientific, USA) plus 2-mercaptoethanol (1% v/v) (Sigma-Aldrich, USA) for each purification procedure, and shaken vigorously using the BeadBug™ Microtube Homogenizer apparatus (D1030-E, Benchmark Scientific). Subsequently, total RNA was isolated using the PureLink RNA Mini kit (ThermoFisher Scientific, USA) in accordance with the manufacturer’s instructions. After isolation, RNA was treated with DNase I (ThermoFisher Scientific). RNA concentration and quality were quantified on a NanoDrop 2000c spectrophotometer (ThermoFisher Scientific) and integrity was evaluated by 2% agarose gel electrophoresis using a UV light photodocumentation system (L-PIX, Loccus Biotecnologia). One microgram of total RNA obtained from RHE was reverse transcribed into complementary DNA using SuperScript™ VILO™ Master Mix, according to the manufacturer’s instructions (ThermoFisher Scientific). Quantitative RT-PCR reactions carried out with a total reaction volume of 10 μL containing 1X of each TaqMan™ designed primers [human; TNF (Hs99999043_m1), IL-1**β** (Hs01555410_m1), IL-6 (Hs00985639_m1), and IL-18 (Hs010003716_m1) ThermoFisher Scientific], TaqMan™ Universal Master Mix II, with UNG (ThermoFisher Scientific), and 10 ng of cDNA resuspended in UltraPure DNase/RNase-free distilled water. No reverse transcriptase controls and template-negative controls were inserted into each assay. The reactions were amplified in a StepOnePlus™ Real-Time PCR Systems thermocycler (ThermoFisher Scientific) under standard conditions. Thermal cycling conditions comprised an initial incubation at 50°C for 2 minutes, 95°C for 10 minutes, 40 cycles of denaturation at 95°C for 15 sec, and annealing and extension at 60°C for 1 min. The relative expression of target genes was normalized by Glyceraldehyde-3-phosphate dehydrogenase (GAPDH; Hs99999905_m1). qPCR data analysis was realized with the N0 method implemented in LinRegPCR v. 2020.2. The results were obtained from 3 independent experiments containing 4 technical replicates, and, for the assay, we performed duplicates for each sample. For the analysis, the mean of data from each experiment was obtained and normalized by the mean of the housekeeping (GAPDH) from each experiment.

### ELISA for TNF-α

For measuring the levels of TNF- α release in the RHE supernatant after 2 or 24 hours of 10 μM oligomeric α-Syn challenge, we used the Human TNF-alpha quantikine ELISA kit (DTA00D, R&D systems) in accordance with the manufacturer’s instructions. Each sample was a pool of at least 3 technical replicates from 3 independent experiments, and for the assay, we performed duplicates for each sample. Immediately after thawing, the samples were centrifuged at 600 x g for 5 minutes to remove particulates. We added 50 μL of assay diluent to each well followed by 50 μL of standard or sample per well and incubated for 2 hours at room temperature on a horizontal orbital microplate shaker. After four washes with a wash buffer, we added 200 μL of Human TNF-α Conjugate to each well and incubated for 2 hours at room temperature on the shaker. Then, we added 200 μL of Human TNF-α conjugate to each well and incubated it for 2 hours at room temperature on the shaker. After repeating the four washes, we added 200 μL of substrate solution to each well and incubated it for 30 minutes at room temperature protected from light. Finally, we added 50 μL of stop solution to each well. Absorbance at 450 nm with wavelength correction set to 540 nm was measured with a Tecan Infinite^®^ 200 PRO (Life Sciences, Switzerland) spectrophotometer.

### Statistical analysis

Data were analyzed either by student-t-test, one-way ANOVA, or two-way ANOVA, followed by the Dunnett test, as applicable. A 95% confidence interval was accepted as statistically significant. All analyses were performed by GraphPad Prism software 8.0.

## Results

### RHE expresses α-Syn, Aβ, and Amyloid precursor protein

Studies have shown that α-Syn, particularly in melanocytes and sensory neurons (Rodriguez-Leyva et al., 2017), as well as Aβ (Heinonen et al., 1994) are found in samples from human skin biopsies. We asked whether the RHE also expresses **α**-Syn, Aβ, and the Amyloid precursor protein (APP), whose proteolysis generates Aβ, as seen in skin biopsies. So, we immunostained RHE for **α**-Syn, Aβ, and APP. We found **α**-Syn immunoreactivity in a few cells from the stratum basale (Figure 1A and B), a diffuse APP immunoreactivity throughout the epidermis (Figure 1C), and Aβ immunoreactivity mostly in upper layers of the RHE (Figure 1D).

**Figure 1:**
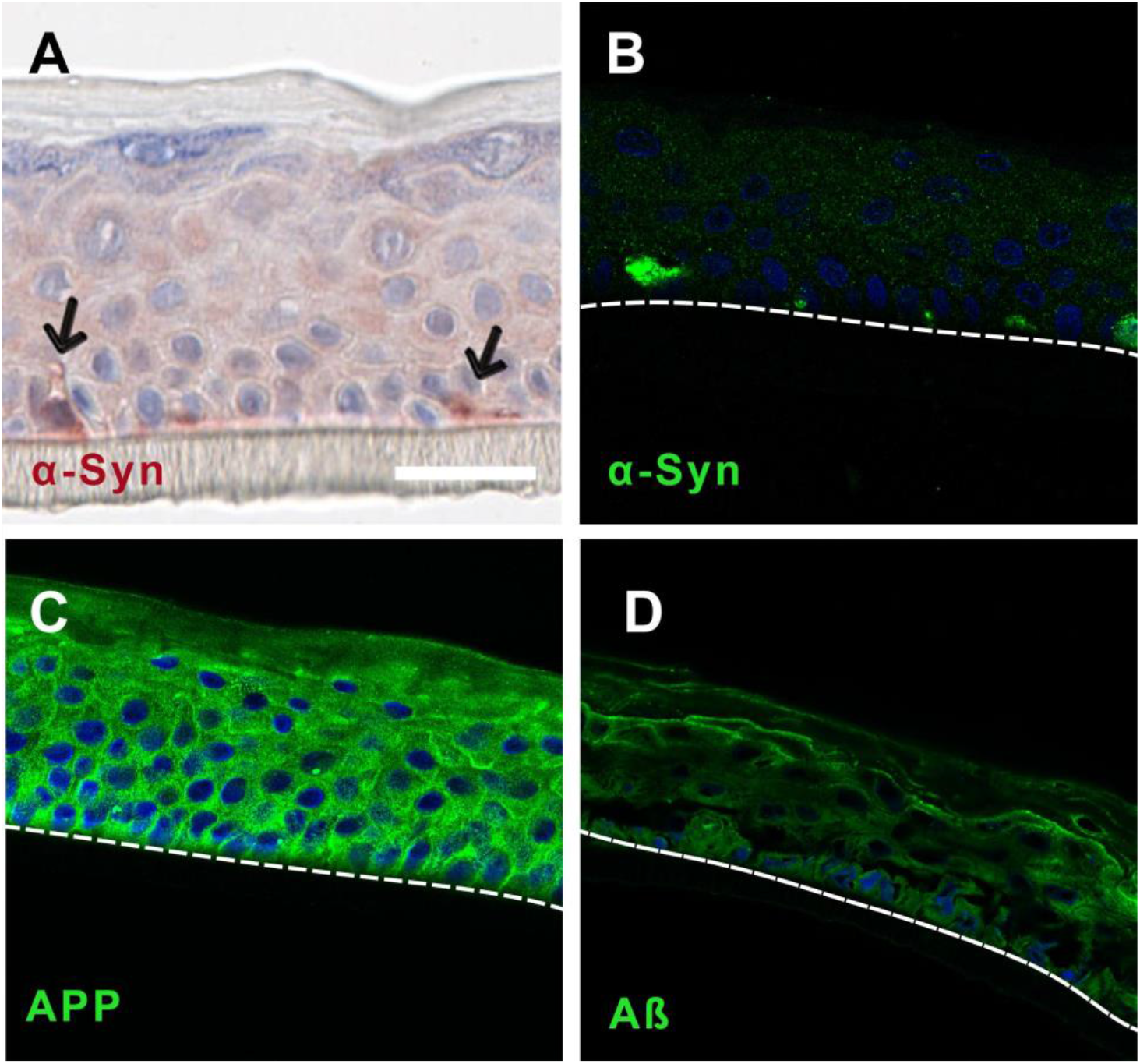
RHE expresses amyloidogenic proteins. Photomicrographs of 17-day-old RHEs immunostained for α-Synuclein (arrow) counterstained with (A) Hematoxylin-Eosin-Safran or (B) DAPI; and for (C) APP and (D) Aβ nuclear stained with DAPI. (B-D) Dashed lines delimitate the bottom of the stratum basale of the epidermis, representing the dermal-epidermal junction. Scale bar: 50 μm.

### Oα-Syn decreases RHE proliferation but does not affect NP neurite outgrowth

Considering that RHE expresses α-Syn, Aβ and APP it is a suitable model to investigate the toxic role of both aggregates α-Syn and Aβ in skin biology. Therefore, we argued whether, under certain conditions, these aggregated proteins would lead to toxic responses. So, we challenged RHE with exogenous 10 μM Oα-Syn or 1 μM aggregated Aβ, or 20 ng/mL TNF-α for 24 hours. We noticed that some treated RHEs were apparently thinner than control ones. We hypothesized that this thinning could be due to an impairment in cell proliferation. Thus, the samples were immunostained for the proliferation marker Ki67, and we confirmed that TNF-α-challenged RHE (2.18 ± 0.54%, p<0.05) and, more pronouncedly, Oα-Syn-challenged RHE (0.72 ± 0.3%, p<0.0001) presented a lower percentage of Ki67 positive nuclei compared with control (5.95 ± 0.71%), and Aβ-challenged RHE (7.75 ± 1.5%) (Figure 2A-D, and E). We asked whether this decrease in proliferation could reflect in RHE thinning. Indeed, Oα-Syn-challenged RHE, but not TNF-α-challenged RHE (58.61 ± 0.66), had a thinner proliferative stratum (55.02 ± 3.53 μm) compared with control RHE (67.5 ± 2.58 μm, p<0.01) (Figure 2F).

**Figure 2:**
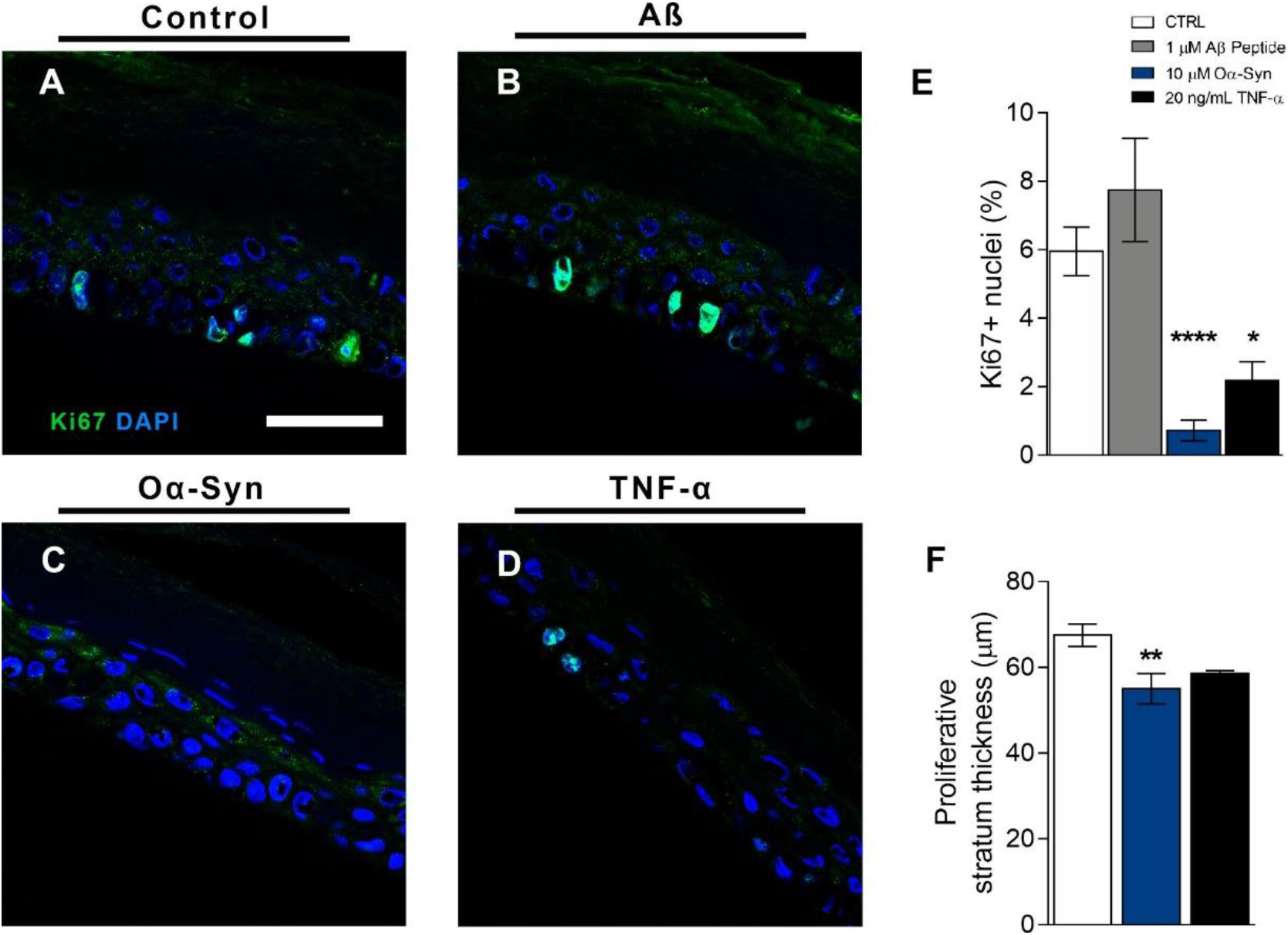
Oα-Syn and TNF-α decreased keratinocyte proliferation. (A-D) Photomicrographs of control and RHE challenged with 10 μM Oα -Syn, 1 μM Aβ, or 20 ng/mL TNF-α for 24 hours, and immunostained for Ki67 (green) and counterstained with DAPI (blue). (E) Percentage of Ki67 + nuclei relative to total nuclei (n=3 experiments, plotted as average ± SEM, analysed by one-way ANOVA followed by Dunnett’ post-hoc test; * *P*< 0.05, **** *P*< 0.0001). (F) Measurement of the thickness of the RHE proliferative stratum (n=3 experiments, plotted as average ± SEM, analyzed by one-way ANOVA followed by Dunnett’ post-hoc test; ** *P*< 0.01). Scale bar: 50 μm.

As skin innervation decreases with age, we asked whether Oα-Syn, which is an aging and neurodegeneration-related protein, could impair human neurite outgrowth. Therefore, we challenged NP with 30 μM Oα-Syn for 24 hours and counted the number of branches crossing a circular mask created at 4 times the diameter of NP’s core, which is composed of nuclei. We observed that Oα-Syn did not affect neurite outgrowth (Control: 43.11 ± 2.94, and Oα-Syn: 43 ± 2.41) (Supplementary figure 1).

**Supplementary figure 1:**
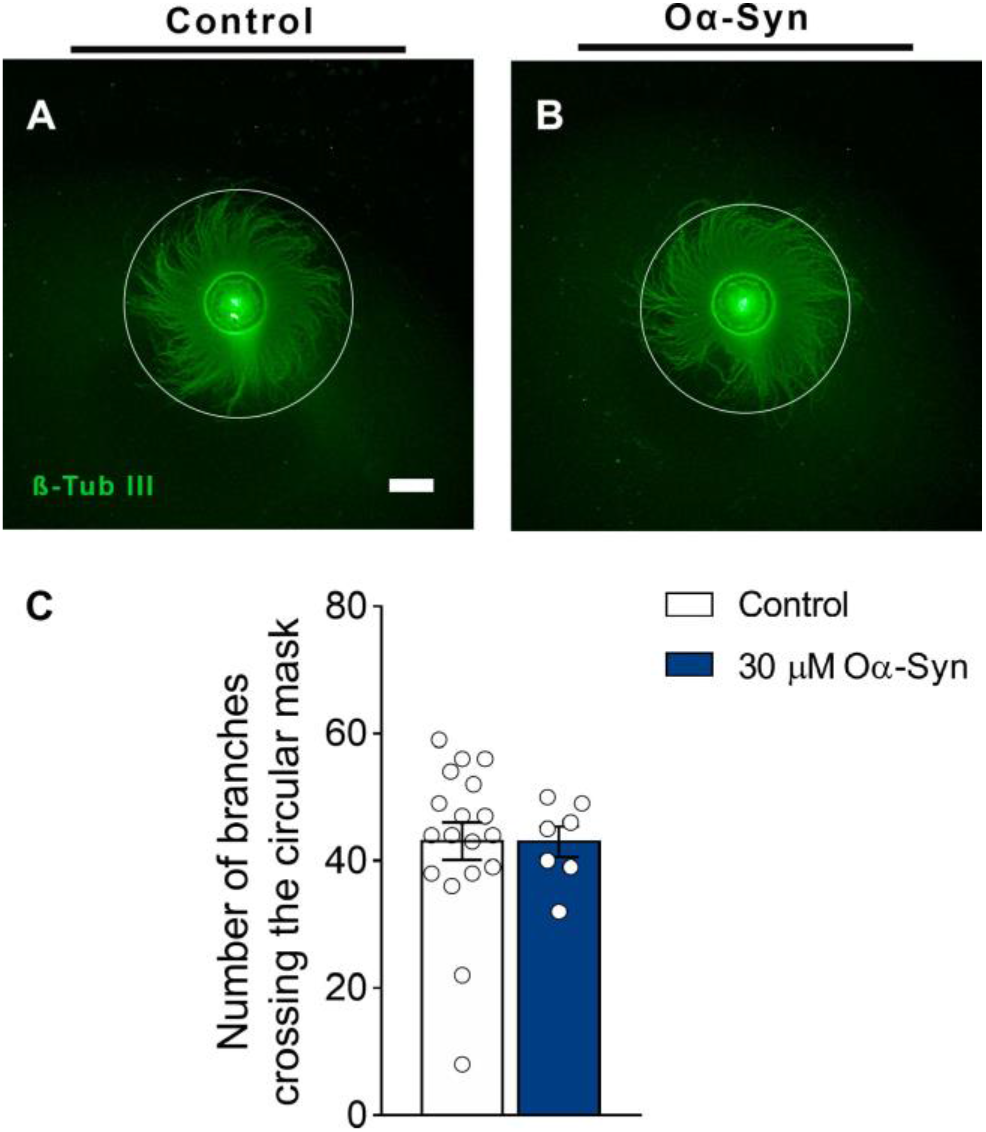
Oα-Syn did not affect neurite outgrowth. (A and B) Photomicrographs of control and NP challenged with 30 μM Oα-Syn for 24 hours and immunostained for β-Tub III. The circular mask at 4 times the diameter of NP’s core was drawn to illustrate the points where neurites cross it. (C) Number of branches crossing the circular mask (n=1 experiment), plotted as average ± SEM, analyzed by t-test. Scale bar: 500 μm.

This anti-proliferative effect of Oα-Syn on the RHE was also observed in 2D culture of HEKn challenged for 24 hours, and immunostained with Ki67 (Figure 3). More importantly, to certify that the oligomeric α-Syn species, which is known as toxic, accounted for the decrease in keratinocyte proliferation, we compared monomeric or oligomeric α-Syn species in HEKn. As expected, we observed that the percentage of Ki67 positive cells was lower in Oα-Syn-challenged HEKn at both 10 μM (56.5 ± 25.66%, p<0.0001) and 30 μM (44.81 ± 37.01%, p<0.0001) concentrations, but not different from monomeric α-Syn-challenged HEKn (96.32 ± 7.4), when compared to control (Figure 3).

**Figure 3:**
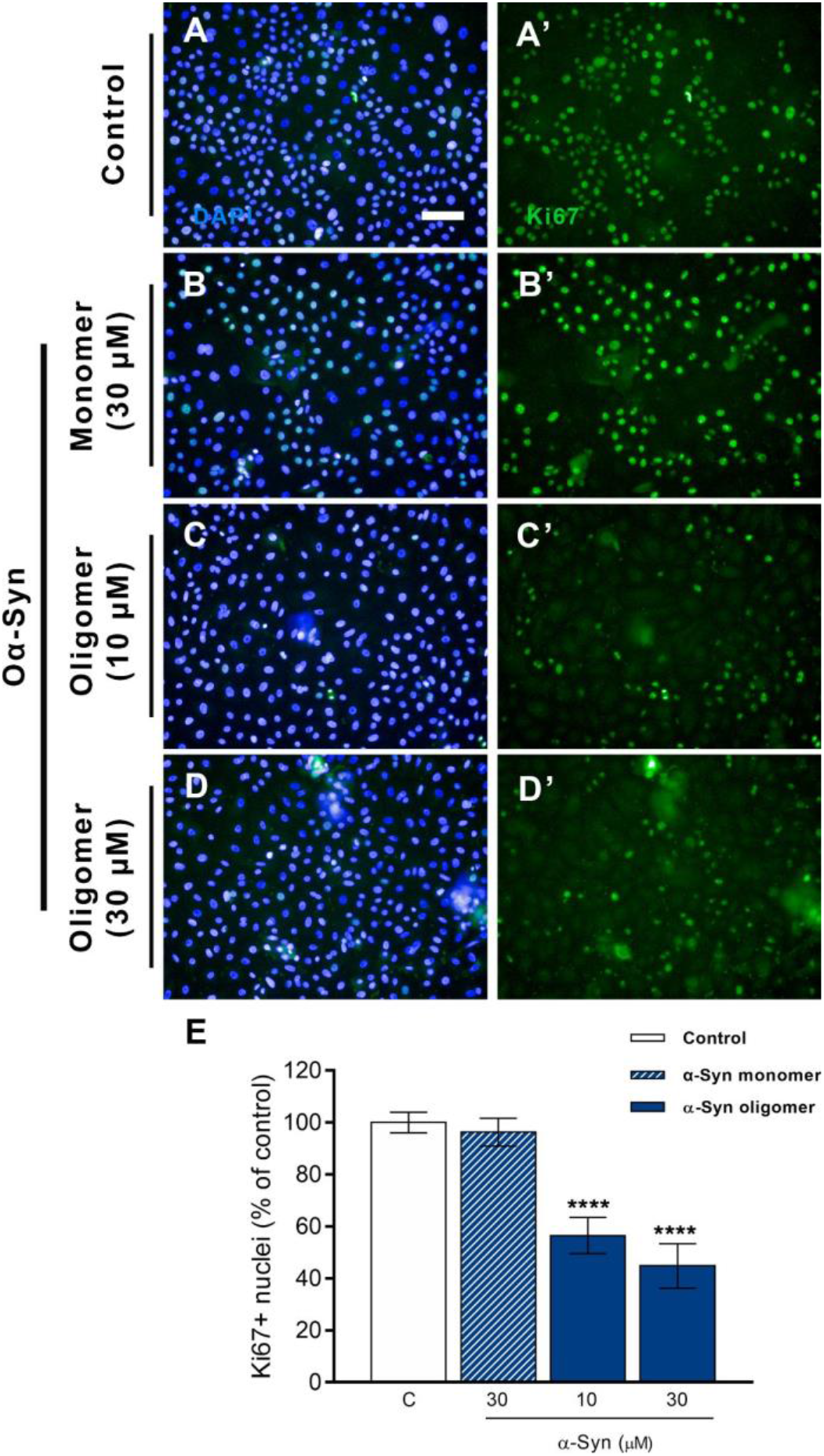
Oligomeric but not monomeric α-Syn accounts for the decrease in proliferation in HEKn. (A-D’) Photomicrographs of control HEKn and challenged with oligomeric (10 μM or 30 μM) or monomeric (30 μM) α-Syn for 24 hours. (A’, B’, C’ and D’) immunostained for Ki67 and (A, B, C, and D), respectively, and nuclei stained with DAPI. (E) Percentage of Ki67 + cells relative to control (n=3 experiments, plotted as average ± SEM, analyzed by one-way ANOVA followed by Dunnett’ post-hoc test; **** *P*< 0.0001). Scale bar: 50 μm.

As Oα-Syn is a toxic protein aggregate that decreases keratinocyte proliferation, we asked whether it may also cause cell death 24 hours following the insult. Then, we counted the total nuclei of 10 μM Oα-Syn-challenged and control HEK. We observed that the number of nuclei from Oα-Syn-challenged HEKn (91.5 ± 4.86%) was comparable with control (100 ± 4.2%) (Supplementary figure 2).

**Supplementary figure 2:**
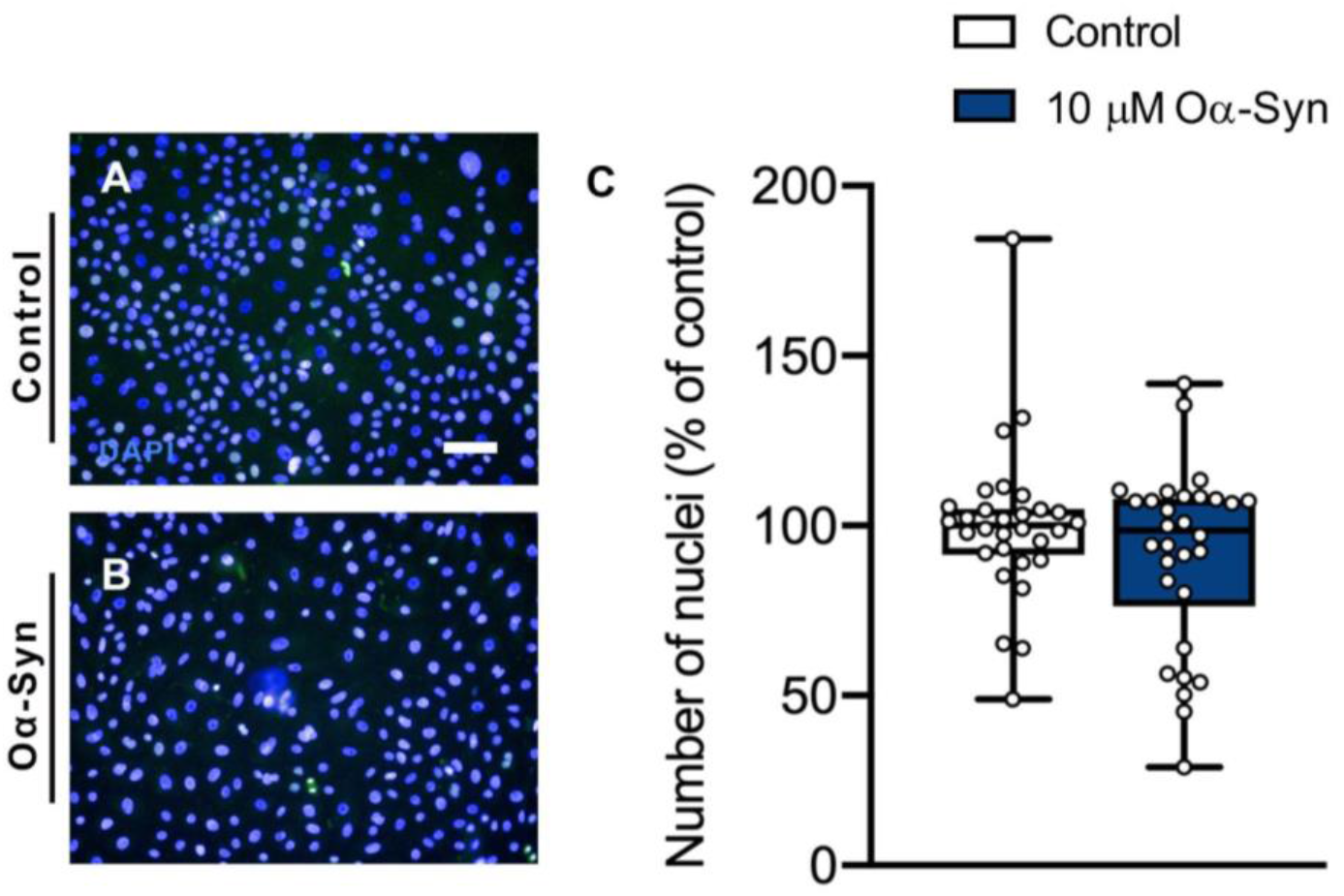
Oα-Syn does not decrease the number of HEKn. (A and B) Photomicrographs of control and HEKn challenged with Oα-Syn for 24 hours stained with DAPI (blue). (C) Percentage of cells relative to control (n=3 experiments with 10 replicates each, plotted as box and whiskers with all individual points, analyzed by unpaired t-test with Welch correction; P= 0.1933). Scale bar: 50 μm.

### Oα-Syn prompts NF-kB nuclear translocation in keratinocytes

TNF-α is a crucial cytokine in skin inflammation and it is induced by stimuli such as UVB (Bashir et al., 2009). TNF-α triggers RelA/p65 NF-kB subunit translocation to the nucleus, thereby regulating various processes including cell proliferation and expression of inflammatory proteins (Joyce et al., 2001; Pupe et al., 2003; Lan et al., 2005). NF-kB nuclear translocation is considered key for the aging process, and several senescence pathways converge through NF-kB (Wang et al., 2019). We sought to verify whether Oα-Syn would also mobilize this pathway. First, we established the NF-kB nuclear translocation response in HEKn and RHE. For this, we challenged HEKn with 2, 10, and 20 ng/mL TNF-α for 45 minutes and observed that TNF-α increased the nuclear/cytoplasm ratio of NF-kB fluorescence intensity at all tested concentrations (1.69 ± 0.03, p< 0.001; 1.64 ± 0.04, p< 0.05; and 1.68 ± 0.02, p<0.001, respectively) compared with control (1.29 ± 0.07) (Supplementary figure 4). Because the RHE is a tridimensional structure composed of layers, we challenged it for 45 minutes only with the highest concentration that induced the translocation response in HEKn, 20 ng/mL, applied from the stratum basale side. We found that this insult also increased the nucleus/cytoplasm NF-kB fluorescence intensity in RHE compared with control (0.94 ± 0.02 and 0.63 ± 0.03, respectively, p< 0.001) (Supplementary figure 3).

**Supplementary Figure 3:**
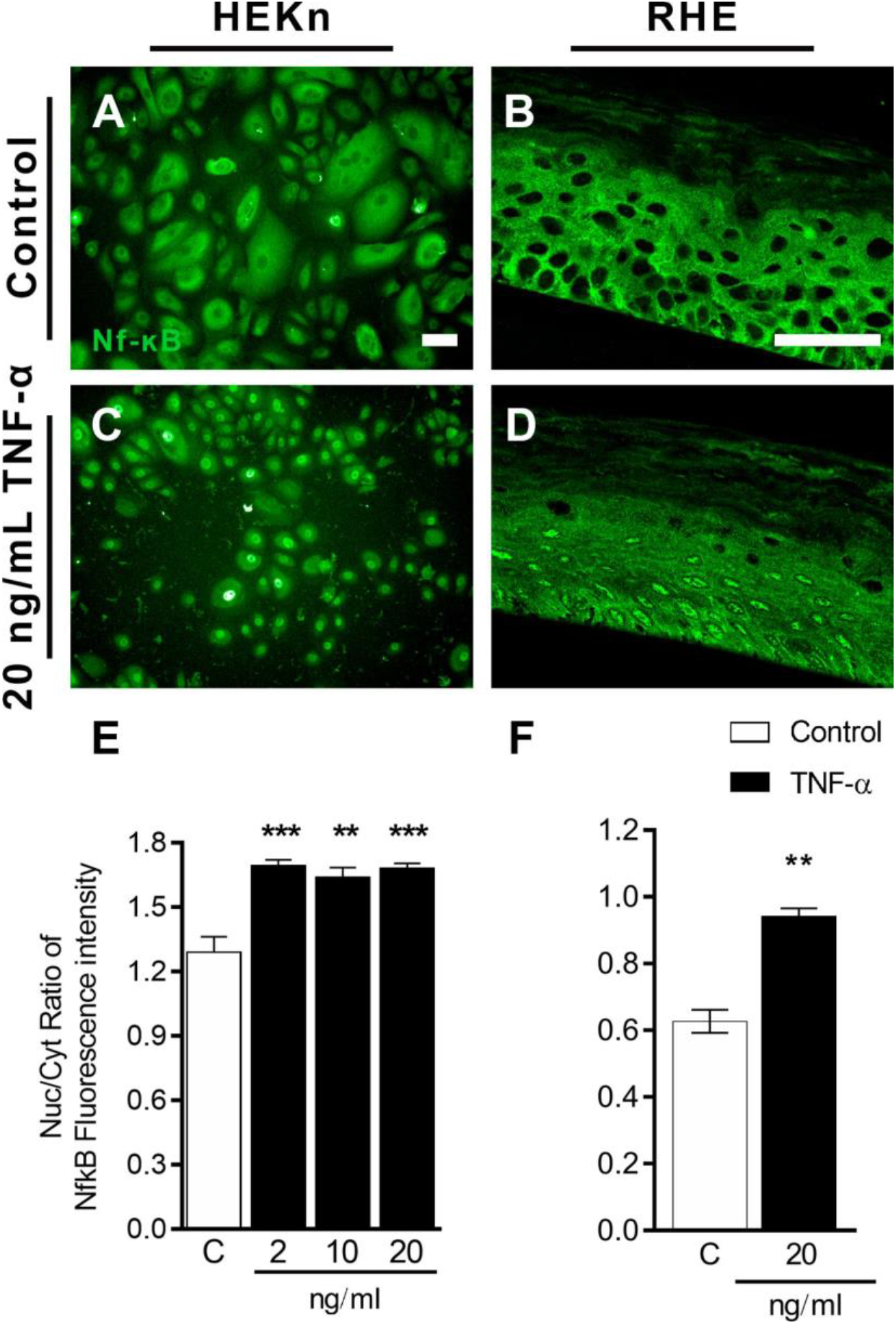
TNF-α insult for 45 minutes triggers NF-kB nuclear translocation in HEKn and RHE. Photomicrographs of control and TNF-α-challenged (A and C) HEKn and (B and D) RHE immunostained for NF-kB. (E and F) Nucleus/cytoplasm NF-kB fluorescence intensity (n=3 experiments, plotted as average ± SEM, analysed by one-way ANOVA followed by Dunnett’ post-hoc test (E) or unpaired t-test; ** *P*< 0.01, *** *P*< 0.001). Scale bar: 50 μm.

After confirming that keratinocytes exhibit the NF-kB nuclear translocation response following insult with the positive control, TNF-α, we investigated the putative inflammatory potential of Oα-Syn on these cells. As Oα-Syn causes inflammation in the nervous system, we wondered if it could have decreased keratinocyte proliferation by eliciting an inflammatory response. Thus, we challenged HEKn with 10 μM Oα-Syn or 20 ng/mL TNF-α for 45 minutes. We found that Oα-Syn-challenged HEKn (1.35 ± 0, p<0.0001) had a higher nuclear/cytoplasm NF-kB fluorescence intensity, equivalent to the positive control (1.28 ± 0.01, p<0.0001), compared with control (1.06 ± 0) (Figure 3).

**Figure 3:**
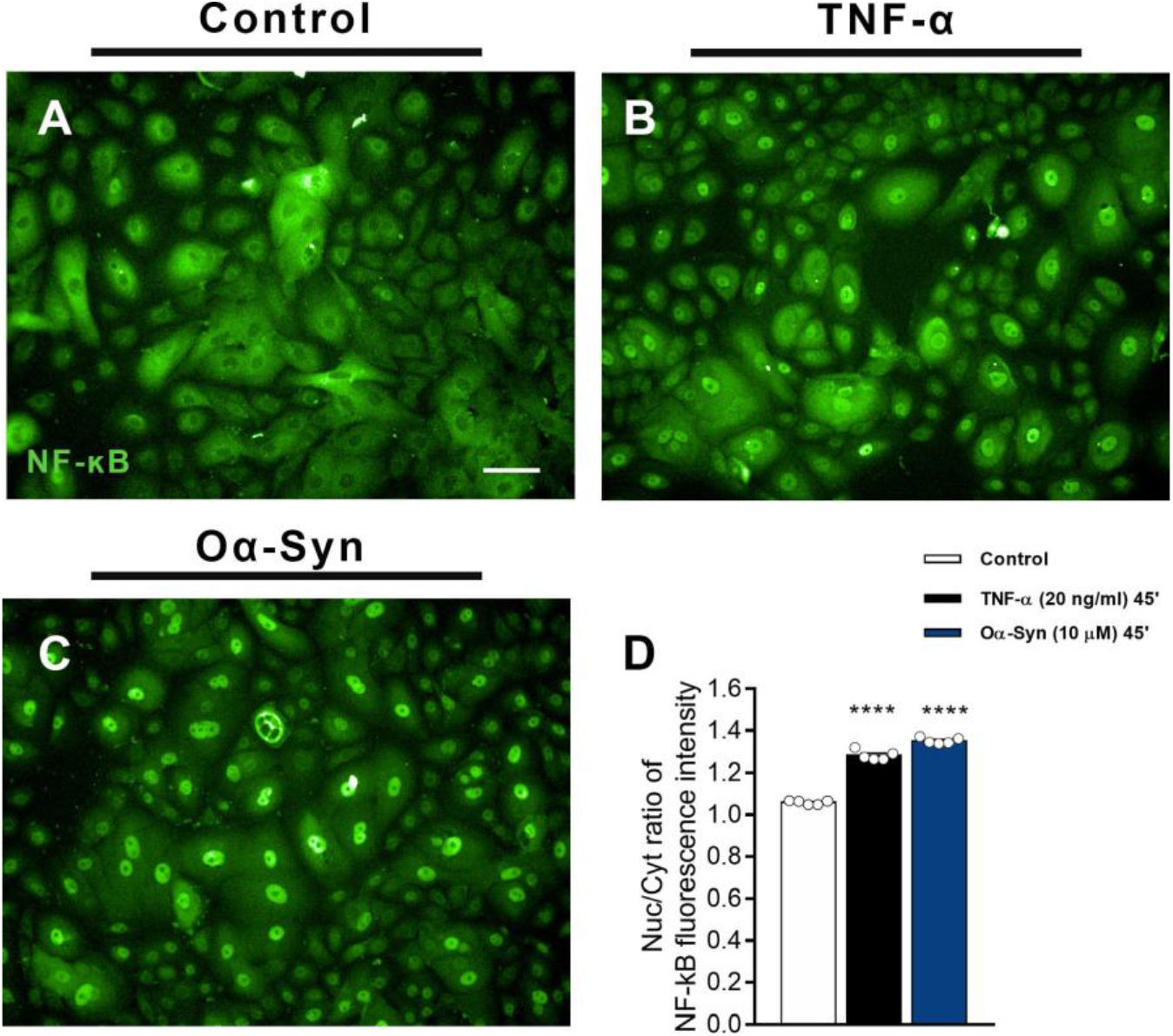
Oα-Syn insult for 45 minutes triggers NF-kB nuclear translocation in HEKn. (A-C) Photomicrographs of control and TNF-α- and Oα-Syn-challenged HEKn immunostained for NF-kB. (D) Nucleus/cytoplasm NF-kB fluorescence intensity (n=2 experiments, plotted as average ± SEM, analyzed by one-way ANOVA followed by Dunnett’ post-hoc test; **** *P*< 0.0001). Scale bar: 50 μm.

### Oα-Syn increases cytokine expression and release in RHE

As Oα-Syn induced NF-kB nuclear translocation, which is a transcription factor involved in the regulation of several inflammatory genes, we asked whether Oα-Syn could increase the mRNA levels of some proinflammatory cytokines in the RHE. We found that 2 hours after the insult, Oα-Syn-challenged RHE presented an increase in IL-1β (Control: 0; Oα-Syn: 0.4, p<0.05) and TNF-α (Control: 0; Oα-Syn: 0.18 ± 0.03, p<0.05) mRNA levels compared with control (Figure 4B and 4C). Other tested cytokines did not significantly change their mRNA levels, such as IL-18 (Control: 0.1 ± 0.03; Oα-Syn: 0.06 ± 0.02) and IL-6 (Control: 0.58 ± 7.35; Oα-Syn: 0) (Figure 4A and 4D). Since TNF-α had a similar effect to Oα-Syn in decreasing keratinocyte proliferation, we figured it was important to also verify the protein release. Indeed, TNF-α release increased following Oα-Syn insult in a time-dependent manner (Oα-Syn 2h versus Oα-Syn 24h, p<0.01) compared with control (Control 2h: 0; Oα-Syn 2h: 864.3 ± 172.75 pg/mL, p<0.05; Control 24h: 7.2 ± 3.74 pg/mL; Oα-Syn 24h: 2,215 ± 264.52 pg/mL, p<0.0001) (Figure 4E).

**Figure 4:**
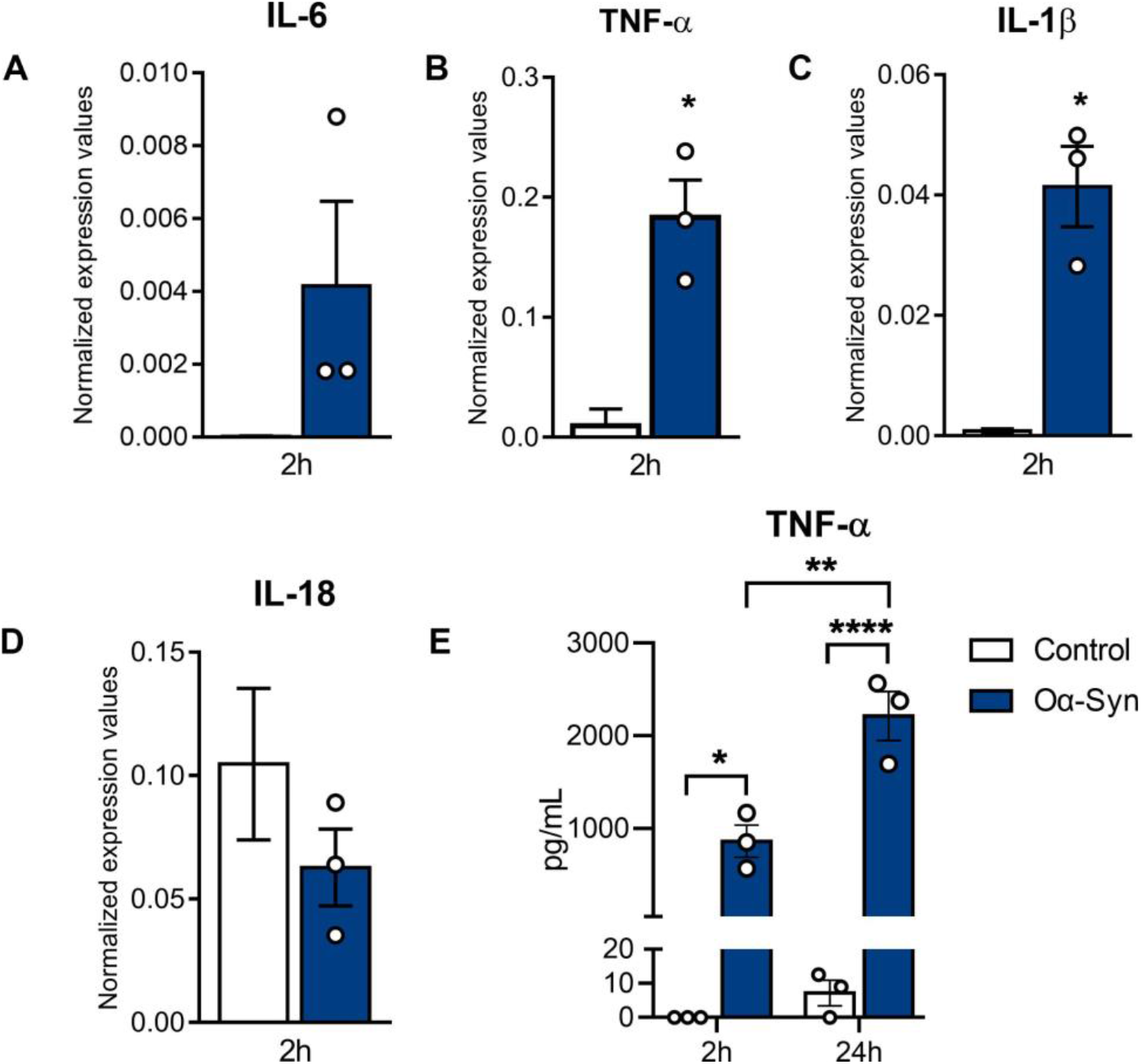
Oα-Syn triggers TNF-α and IL-1β overexpression and the release of TNF-α by the RHE. (A-D) RT-qPCR for the proinflammatory cytokines IL-6, TNF-α, IL-1β, and IL-18 2 hours after challenge, plotted as absolute values normalized by the housekeeping (GAPDH). (E) ELISA for TNF-α released in the RHE supernatant at 2 and 24 hours after the challenge. n=3 experiments, plotted as average ± SEM, analysed by paired t-test (A-D) or two-way ANOVA followed by Tukey’s post-hoc test (E); * *P*< 0.05, ** *P*< 0.01, **** *P*< 0.0001).

## Discussion

The aggregation of proteins and peptides, such as α-Syn, Aβ, and tau, is one of the hallmarks of age-related neuroinflammation (Glass et al., 2010). However, a growing body of evidence supports the presence of these aggregates in tissues outside the central nervous system. For instance, both monomeric and oligomeric α-Syn were found in the plasma, cerebrospinal fluid (Park et al., 2011), and basal tears from patients with Parkison’s disease but also healthy elderly (Hamm-Alvarez et al., 2019). Phosphorylated-α-Syn, which is a post-transductional modification of α-Syn promoting toxic aggregation, was found in postmortem skin biopsies from individuals with non-synucleinopathies neurodegenerative diseases and from one non-neurodegenerative control (Wang et al., 2021). In addition, α-Syn expression was observed in the skin of patients with α-synucleinopathies as well as healthy individuals (Rodriguez-Leyva et al., 2016). Using spectroscopy of human skin biopsy samples, Akerman and collaborators (2019) showed that some amyloidogenic proteins, including α-Syn, build up as the skin ages (Akerman et al., 2019a). Even though it is not an aging model, we also found Aβ, APP, and α-Syn expression in the RHE, the latter being restricted to a few cells from the stratum basale. The expression of α-Syn in the skin has been mainly ascribed to peripheral nerve terminals and melanocytes (Rodriguez-Leyva et al., 2016; Kim et al., 2019). Dermal fibroblasts are also another possible source of α-Syn accumulation in the aged skin, releasing exosomes containing α-Syn and pro-inflammatory mediators by cell-to-cell transmission from the dermis to the epidermis (Danzer et al., 2012; Cerri et al., 2021). However, one study also showed α-Syn immunopositive keratinocytes from the skin of patients with Parkinson’s disease and atypical parkinsonism (Rodríguez-Leyva et al., 2014). One hypothesis is that the melanocytes, which can seldom develop in the RHE, expressed the α-Syn observed in our study, but one cannot rule out the possibility that scarce basal keratinocytes produce it. Regardless of the source, both α-Syn species might coexist, as seen in other tissues, and aggregation of monomeric α-Syn in the skin might occur at some level as the skin ages.

The evidence of α-Syn, Aβ, and APP expression in the skin, along with our findings in the RHE, prompted us to explore the role of these aggregates on RHE physiology. Although other groups have shown a potential role of APP in the epidermis, serving as an epidermal growth factor stimulating keratinocytes proliferation and migration (Herzog et al., 2004; Siemes et al., 2006), Aβ did not affect RHE in our experimental conditions. On the other hand, Oα-Syn caused a decrease in epidermal thickness, which is similar to what happens in senior skin as a result of a decrease in cellular proliferation and reduced innervation (Besné et al., 2002; Gilhar et al., 2004). Interestingly, both TNF-α and, more pronouncedly, Oα-Syn, caused a decrease in the cell proliferation of the stratum basale. TNF-α represses the transcription of genes implicated in the cell cycle in human keratinocytes (Pillai et al., 1989; Kono et al., 1990; Banno et al., 2004), and inhibits human keratinocyte proliferation *in vitro* in a time- and concentration-dependent manner (Detmar and Orfanos, 1990). With regard to Oα-Syn, this is the first evidence that this aggregate directly impacts cell proliferation. However, Oα-Syn plays a notable role in neuroinflammation, causing, for instance, the release of pro-inflammatory cytokines including TNF-α from microglia *in vitro* (Roodveldt et al., 2010). In line with this work, Oα-Syn also increased TNF-α expression and release from RHE. Since this cytokine has known antiproliferative effects in the skin, it is possible that Oα-Syn acted through TNF-α release to inhibit keratinocyte proliferation.

In addition to acting as a toxic protein in the RHE, Oα-Syn is well-known as neurotoxic. The epidermis is innervated by peripheral nervous endings, whose density is affected by aging (Besné et al., 2002). Studies have shown that Oα-Syn increases dopaminergic neuronal reactive oxygen species and apoptosis (Xu et al., 2002), and impairs neurite outgrowth (Koch et al., 2015). However, in this work, Oα-Syn did not affect NP neurite outgrowth. Overexpression of different variants of Oα-Syn acted differently on neurite network morphology in a human dopaminergic neuronal cell line — while E57K Oα-Syn decreased the number of neurites, wild type Oα-Syn did not affect this parameter (Prots et al., 2013). In addition, another work showed that wild type, A30P, and A53T Oα-Syn variants decreased the number of neurites from dopaminergic neurons, but only A30P and A53T did it in non-dopaminergic neurons (Koch et al., 2015). In line with these works, our NP was composed of non-specific neurons that were challenged with wild-type Oα-Syn, which might explain the unaltered neurite morphology observed.

NF-kB is considered the central regulator of inflammatory processes and, not coincidentally, the transcription factor most associated with aging (Tilstra et al., 2011). Both TNF-α and UVB-irradiation induce NF-kB DNA-binding activity in human keratinocytes (Lewis and Spandau, 2007). Here we demonstrated for the first time that Oα-Syn also elicited NF-kB nuclear translocation in keratinocytes with similar intensity as the classical NF-kB inducer, TNF-α. One of the mechanisms by which extracellular Oα-Syn exerts its toxic effects on microglia and astrocytes is through activating Toll-like receptors (TLRs) 1, 2 or 4. This activation, in turn, leads to the nuclear translocation of NF-kB, which triggers the expression of proinflammatory cytokines, such as IL-6, TNF-α, and IL-1β, and induces accumulation of reactive oxygen species and mitochondrial disturbance. Human keratinocytes are known to express TLR1, TLR2, and TLR4 (Pivarcsi et al., 2004; Köllisch et al., 2005). Herein, we found that exogenous Oα-Syn caused the nuclear translocation of NF-kB and overexpression of proinflammatory cytokines. Therefore, TLR is the most likely pathway involved in the Oα-Syn-mediated toxic effects observed in keratinocytes.

Common *in vitro* models used to induce skin stress and premature senescence are hydrogen peroxide insult or UVB irradiation. Besides triggering the release of reactive oxygen species, these stimuli also trigger the release of inflammatory signals, including TNF-α and IL-1β (Bashir et al., 2009; Rinnerthaler et al., 2015; Song et al., 2018). In this work, Oα-Syn also induced RHE inflammation by increasing TNF-α and IL-1β expression. The latter is a cytokine that is released by activation of the NALP3 inflammasome, which has also been implicated in Oα-Syn-induced responses by microglia (Trudler et al., 2021). Both cytokines IL-1β and TNF-α overexpression may be triggering positive feedback of inflammatory cytokines previously described in skin inflammaging studies (Grover and Grewal, 2008; Zhuang and Lyga, 2014; Fuller, 2019; Pilkington et al., 2021) resulting in keratinocyte loss, epidermal degeneration, and ultimately, fine lines and impaired mechanic skin properties. Aging skin structural changes include reduced elasticity, as well as reduced numbers and altered morphology of tactile receptors, consequently impairing skin sensitivity and sensoriality in the elderly (McIntyre et al., 2021).

α-Syn is poorly expressed in the RHE and, as a prion-like protein, it can be transported by cell-to-cell propagation. Accordingly, our hypothesis is that peripheral nerve terminals and melanocytes, which are the main cells expressing α-Syn in the skin, convey α-Syn to the epidermis. Interestingly, α-Syn is found in postmortem tissues of the peripheral nervous system from patients with α-synucleinopathy (Sumikura et al., 2015), in sensory nerve terminals in the skin (Akerman et al., 2019a), and in sensory neurons derived from human induced pluripotent stem cells (Guimarães et al., 2018). Therefore, simply by accumulation or via other factors, such as UV radiation (Carmo-Gonçalves et al., 2014), α-Syn aggregates into Oα-Syn, causing skin inflammation and degeneration (Figure 5). Even though there are few reports that distinguish Oα-Syn from its monomeric form in healthy individuals, there is evidence that even picomolar concentrations of the oligomers mixed with the more abundant monomers are sufficient to elicit TNF-α release from microglia (Hughes et al., 2019). In contrast, in the same work, Oα-Syn alone had an EC50 of around 10 μM in causing TNF-α release, a concentration consistent with the ones used in the current work.

**Figure 5:**
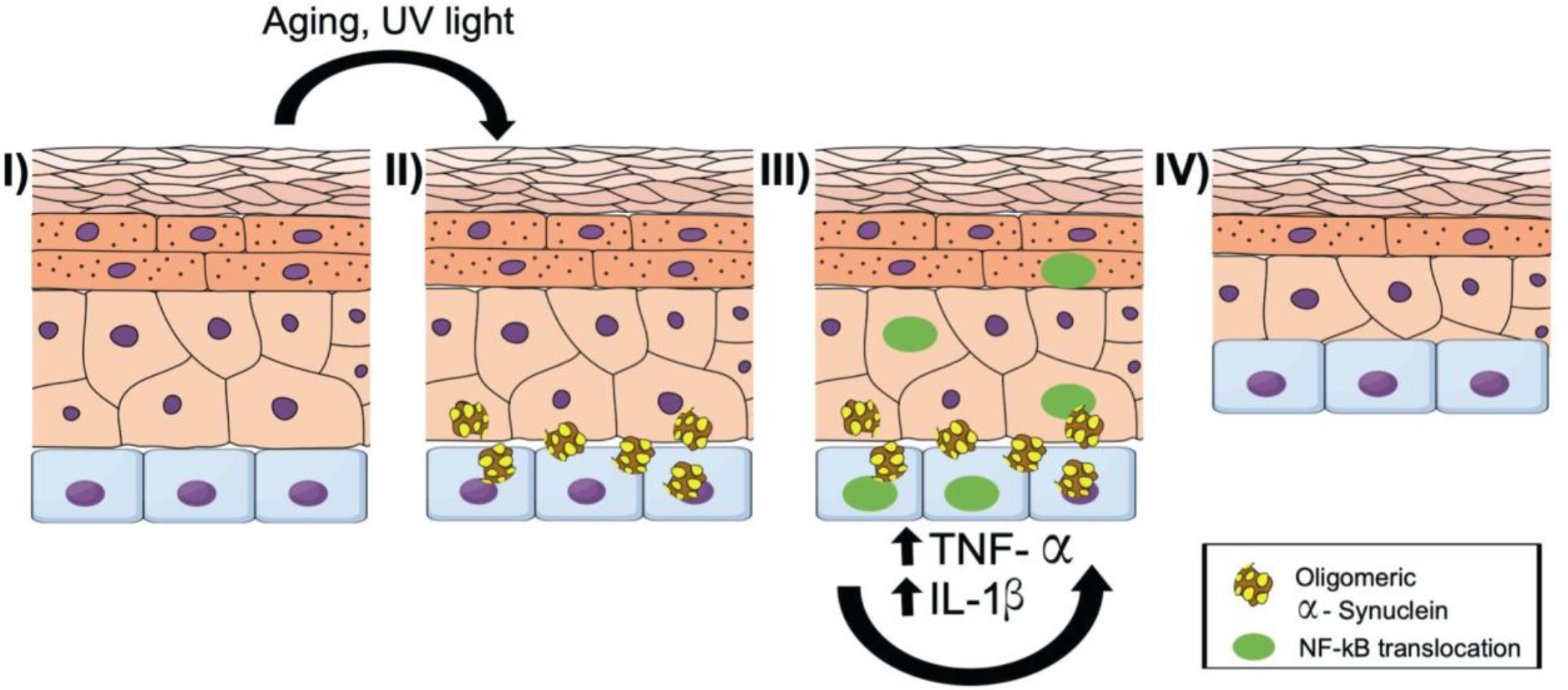
The working hypothesis for the role of Oα-Syn in skin degeneration with aging. I) UV exposure and aging might contribute to oligomerization of available α-Syn. II) α-Syn aggregates and accumulates in the epidermis. III) Oα-Syn activates inflammatory responses in keratinocytes, via NF-kB translocation to the nucleus (represented by green nuclei), and triggers the transcription and autocrine release of inflammatory cytokines, such as TNF-α and IL-1β. IV) One of the observed effects of these cytokines on the epidermis is the reduced keratinocyte proliferation, which contributes to skin degeneration and, ultimately, to epidermal thinning, characteristic of aged skin. Illustration prepared with Mind the Graph (https://mindthegraph.com).

Oα-Syn remarkably affected the epidermal basal layer, which is critical to tissue regeneration because it mostly contains proliferative keratinocytes. Decreased proliferation caused by Oα-Syn yielded a thin epidermis as seen in sun-damaged, aged, as well as diseased skin. Our translational model allows the understanding of aspects of skin aging, observed in clinical studies related to neurodegenerative diseases, but also important on skin homeostasis in aging. This study opens up new avenues for the searching of therapies for α-Syn accumulation in the skin, important not only for skin degeneration caused by neurodegenerative diseases but also for normal aging.

